# Covariant Biochemical Systems Theory: cBST1–cBST3 Descriptors and Quantitative Validation

**DOI:** 10.64898/2026.07.26.732504

**Authors:** Chikoo Oosawa

## Abstract

Biochemical Systems Theory (BST) represents nonlinear biochemical rate laws by local power-law approximations in logarithmic concentration coordinates. First-order coefficients are elasticities, whereas higher-order derivatives describe local log-synergism and its variation. Ordinary higher derivatives, however, are not tensorial under nonlinear reparameterizations and can mix biochemical response structure with coordinate artifacts. We formulate a covariant hierarchy, cBST1–cBST3, on a positive operating-point space equipped with a declared reference connection. cBST1 recovers classical elasticities in a flat logarithmic chart, cBST2 is the covariant Hessian of the log-response, and cBST3 is the symmetrized covariant derivative of cBST2. The framework is quantitatively evaluated using three representative rate laws from the curated yeast glycolysis model BIOMD0000000064: glucose transport, glucose phosphorylation, and phosphofructokinase. For 10,000 finite log-concentration perturbations at each of five radii, cBST2 reduced the cBST1 log-rate root-mean-square error by 96.6–98.6% at the largest tested radius, and cBST3 provided a further 68.4–98.1% reduction. The contracted cBST2 and cBST3 terms strongly predicted the corresponding lower-order truncation errors. Under the nonlinear transformation *q_i_* = sinh(*u_i_*), covariant contractions agreed across coordinates to within a 95th-percentile relative error of 1.2 × 10^−13^, whereas ordinary higher derivatives showed order-unity coordinate mismatches. Supplementary analytic tests recovered the expected second-, third-, and fourth-order truncation-error scaling. These results show that cBST1–cBST3 are not only coordinate-consistent descriptors but also practical diagnostics of where local power-law approximations require higher-order correction.

## 1 Introduction

Biochemical Systems Theory (BST) and S-system modeling provide a compact power-law language for representing nonlinear biochemical networks (Savageau, 1969a,b, 1970, 1976; Voit, 2000, 2013; Salvador, 2024). For a positive rate law *v*(*x*), with positive concentrations *x*_*i*_ *>* 0, the logarithmic response

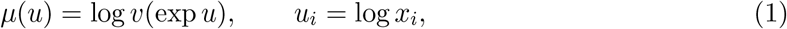

can be approximated locally by a linear function of *u*. Exponentiating this linear approximation gives a multiplicative power law. The linear coefficients are the familiar elasticities or kinetic orders, which quantify local sensitivity of a rate to fractional concentration changes. This viewpoint underlies S-systems, generalized mass-action models, structural kinetic modeling, and local analysis of biochemical networks (Heinrich and Schuster, 1996; Fell, 1997; Steuer et al., 2006; Massing et al., 2022).

The same logarithmic expansion contains higher-order biological information. Second-order extensions of BST were already recognized as a way to increase predictive power and to distinguish mechanisms that may be indistinguishable at first order (Cascante et al., 1991). The Hessian of *µ* in log coordinates measures how elasticities change with the operating point and how two inputs interact nonmultiplicatively. This second-order differential structure is naturally interpreted as a local form of log-synergism: it detects whether the combined fractional effect of two variables differs from what would be predicted by independent local elasticities. This interpretation is closely related to Salvador’s synergism analysis of biochemical systems, including its tensor formulation and treatment of stoichiometric constraints (Salvador, 2000a,b). Third-order terms describe how this synergism varies over the operating region. Such descriptors are useful when comparing kinetic mechanisms, assessing saturation, deciding whether a first-order power-law model is adequate, or transferring local models between different normalizations of the same biochemical system.

A difficulty is that ordinary higher derivatives of a scalar are not themselves coordinate-invariant descriptors. If variables are rescaled affinely in log space, classical BST coefficients transform in a simple way. In practice, however, biochemical models are frequently written in normalized concentrations, saturation variables, transformed thermodynamic forces, latent reduced coordinates, or coarse-grained coordinates that need not be affine functions of log *x*. Under such nonlinear reparameterizations, the ordinary Hessian of a log-response acquires extra terms generated by the coordinate map. If these terms are interpreted directly as biochemical curvature or synergism, a coordinate artifact is mistaken for an intrinsic property of the rate law. This paper formulates a covariant hierarchy of BST descriptors, denoted cBST1–cBST3. The construction is local and geometric, but the intended role of the geometry is modest: it records which coordinate effects are regarded as representational. In this sense, cBST2 should be read as a coordinate-consistent continuation of earlier second-order BST and log-synergism ideas, not as a claim that second-order response analysis is new by itself. A positive operating-point space *M* is equipped with a specified torsion-free reference connection. For each scalar log-response *µ* : *M* → ℝ, cBST1 is the covariant first derivative, cBST2 is the covariant Hessian, and cBST3 is the symmetrized covariant derivative of cBST2. In the flat logarithmic chart used by classical BST, these objects reduce to the ordinary first, second, and third log derivatives. Under nonlinear coordinate changes they transform as tensors, so their contractions with tangent perturbations or fluctuation covariances are coordinate-consistent scalars.

The goal is not to replace classical BST. Instead, the goal is to clarify which parts of BST are intrinsic local descriptors and which parts are tied to a chosen coordinate representation. Classical S-system terms remain exact cBST1 models in the flat log chart. The new role of cBST2 and cBST3 is to quantify, in a coordinate-consistent manner, the local response structure omitted by a single power-law flux. This formulation is also intended to connect BST with structural kinetic modeling, information geometry, and local approximation-risk diagnostics without making these downstream applications prerequisites for the definitions.

A concrete use case is the comparison of enzyme or regulatory responses that have been reported in different local coordinates. The same kinetic mechanism may be expressed in raw concentrations, normalized concentrations, fractional saturation variables, or reduced variables obtained by coarse-graining a larger network. cBST1–cBST3 allow the local exponent, log-synergism, and variation of log-synergism to be compared after the intended geometric structure has been specified. This is particularly useful for distinguishing a true interaction between biochemical inputs from curvature introduced only by a nonlinear coordinate representation.

The remainder of the paper is organized as follows. Section 2 recalls the classical local BST expansion and identifies the coordinate-dependence problem. Section 3 introduces the operating-point geometry and gives practical recommendations for choosing a connection. Section 4 defines cBST1–cBST3, and Section 5 interprets S-system fluxes in cBST form. Section 6 gives analytically controlled response examples. Section 7 describes quantitative tests using representative rates from a curated yeast glycolysis model. Section 8 reports finite-perturbation accuracy, truncation-error diagnostics, and coordinate-consistency results. Section 9 discusses biological interpretation, approximation risk, and limitations.

Figure 1 gives a compact numerical visualization of the first three cBST descriptors for a Hill response. The figure is not needed for the definitions, but it helps distinguish the three roles: cBST1 is the local exponent, cBST2 is the local curvature or log-synergism, and cBST3 is the local variation of that curvature.

**Figure 1.**
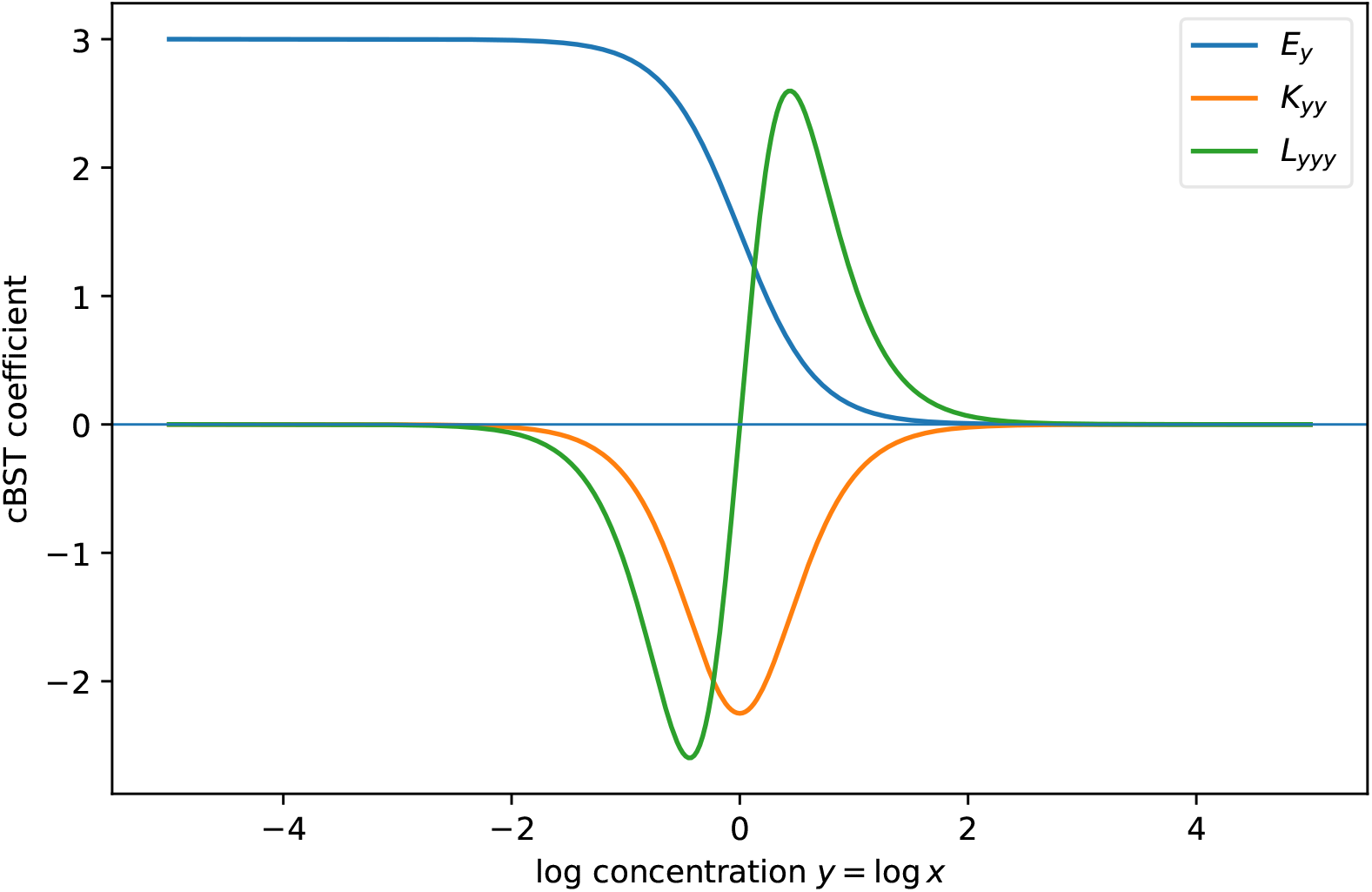
Illustrative cBST descriptors for a one-dimensional Hill activation response with *n* = 3 and *K* = 1 in the flat logarithmic chart. The first-order coefficient tracks the effective local exponent, the second-order coefficient marks the curvature associated with the saturation transition, and the third-order coefficient changes sign across the transition because the curvature is increasing on one side and decreasing on the other.

## 2 Classical local BST expansion

Let 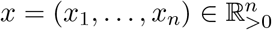 be positive concentrations and let *v*(*x*) *>* 0 be a smooth rate law. In logarithmic coordinates *u*_*i*_ = log *x*_*i*_, define

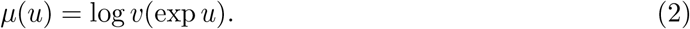

At an operating point *u*_*\**_, the ordinary log-Taylor expansion is

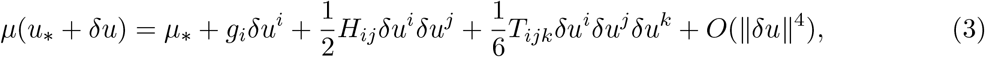

where

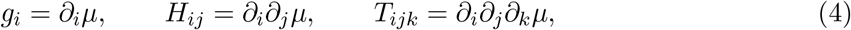

all evaluated at *u*_*\**_. The first-order truncation gives

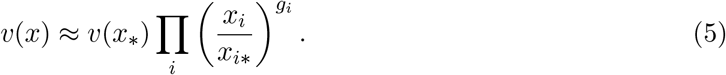

Thus

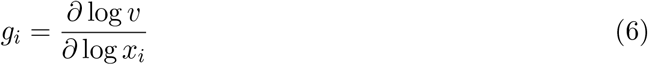

is the local power-law exponent or elasticity. The second-order term *H*_*ij*_ measures the local change of elasticities and the differential form of log-synergism. The third-order tensor *T*_*ijk*_ measures how this local synergism itself changes with the operating point.

Equation (3) is fully appropriate in a fixed affine log chart. The difficulty appears when one changes coordinates nonlinearly. If *u* = *u*(*q*), the ordinary second derivative of the transformed scalar 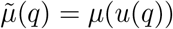 obeys

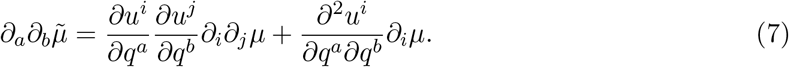

The second term is produced purely by the nonlinear coordinate map. If it is interpreted as biochemical curvature, a coordinate artifact is mistaken for response structure. An analogous problem occurs at third order. This motivates replacing ordinary derivatives by covariant derivatives relative to a specified connection.

## 3 Operating-point space and reference connection

Let *M* be a smooth *n*-dimensional operating-point space whose points represent logarithmic concentrations, reduced biochemical coordinates, or local parameters. A coordinate chart is denoted by *q* = (*q*^1^, …, *q*^*n*^). A scalar log-response is a smooth map

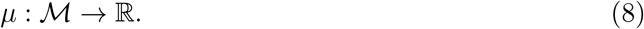

To compare higher derivatives across coordinate systems, equip *M* with a specified torsion-free reference connection ∇. In the present paper, the connection should be read as a modeling convention for comparing local power-law descriptions at nearby operating points. It is not introduced to impose Riemannian geometry on biochemical kinetics. For classical BST, the natural convention is the flat connection in logarithmic concentration coordinates.

The connection is part of the model specification. This point is important for biological interpretation. If the aim is to compare classical BST elasticities and S-system exponents, the flat log connection should be used unless there is a clear reason to choose otherwise. If the variables are nonlinear reduced coordinates derived from a mechanistic transformation, the induced connection should be transformed together with the coordinates. If an independent metric is scientifically meaningful, for example through a statistical observation model or a Fisher–Rao metric, a metric-compatible connection such as the Levi-Civita connection may be considered as an optional choice (Amari and Nagaoka, 2000; Ay et al., 2017; Nielsen, 2020). Such a choice is not the default for classical BST and must be reported as part of the model.

A practical rule is therefore to choose the connection according to what is meant to be held fixed. If the biochemical operating point is being described in ordinary logarithmic concentrations, the flat log connection preserves the classical BST interpretation. If a nonlinear transformation is only a change of chart, the connection should be transformed with the chart so that an exact power law remains exact. If a reduced coordinate is a model-based coarse-graining, the connection encodes the reduction geometry and should be reported with the reduced model. These choices make the framework connection-dependent but not arbitrary: the declared connection specifies which coordinate effects are regarded as representational rather than biochemical.

For a scalar field *µ*, the first covariant derivative equals the ordinary differential,

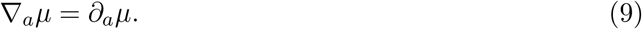

The second covariant derivative is

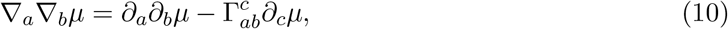

where 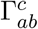 are connection coefficients. The connection term cancels the extra term in Eq. (7) when that term is generated solely by a nonlinear reparameterization. Higher covariant derivatives are formed by applying ∇ to the corresponding tensor fields.

The covariant Taylor interpretation is local. Let *ξ* ∈ *T*_*q\**_ *M* be a small tangent vector and let *q* = exp_*q\**_ (*ξ*) be the geodesic displacement under the selected connection. In normal coordinates at *q*_*\**_, the Christoffel symbols vanish at the base point and the covariant Taylor series has the same algebraic form as the ordinary Taylor series, but its coefficients are tensors. This is the local setting in which the cBST hierarchy is defined.

## 4 The cBST1–cBST3 hierarchy

### Definition 1

(cBST1: covariant elasticity). *For a scalar log-response µ on M, the first-order covariant BST tensor is*

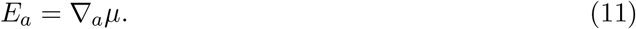

*It is a covector on T* ^*\**^*M and represents coordinate-consistent local elasticity*.

### Definition 2

(cBST2: covariant log-synergism). *The second-order covariant BST tensor is*

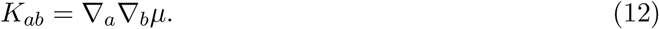

*For a torsion-free connection, K*_*ab*_ = *K*_*ba*_. *It represents coordinate-consistent log-synergism or curvature of the log-response*.

### Definition 3

(cBST3: covariant variation of log-synergism). *The third-order covariant BST tensor is the symmetrized covariant derivative of K*,

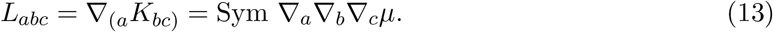

*It represents the local covariant variation of log-synergism*.

### Remark 1

(Why the present study stops at cBST3). *The same construction can formally be continued to any order*,

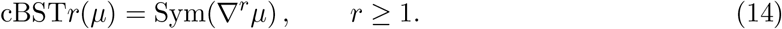

*The present study stops at third order for scientific rather than algebraic reasons. cBST1, cBST2, and cBST3 have direct local interpretations as elasticity, log-synergism, and variation of log-synergism, respectively, and the cubic truncation already leaves an O*(∥*ξ*∥^4^) *local remainder. Higher orders rapidly increase the number of independent components: a symmetric rank-r tensor in n variables has* 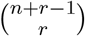 *components, giving 220 components for cBST3 and 715 for cBST4 when n* = 10. *Estimation of cBST4 and above from noisy biochemical measurements would therefore require substantially denser perturbation designs and stronger regularization. Such terms may be useful for exceptionally sharp switches or as residual diagnostics for a cBST3 approximation, but their empirical stability and biological interpretation warrant a separate study*.

The hierarchy gives the local approximation

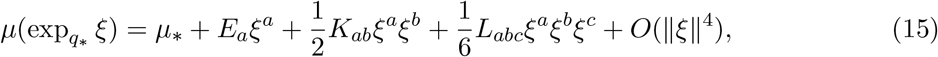

where all tensors are evaluated at *q*_*\**_. Exponentiating the first-order truncation yields the local power-law form in affine log coordinates, while the higher terms quantify departures from a locally exact power law.

### Proposition 1

(Reduction to ordinary BST in a flat log chart). *Suppose M* = ℝ^*n*^ *is equipped with logarithmic coordinates u*_*i*_ = log *x*_*i*_ *and the flat connection for which* 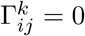 *in the u chart*.

*Then cBST1, cBST2, and cBST3 reduce to the ordinary first, second, and third log derivatives in Eq. (3). In particular, cBST1 gives the classical elasticity vector*.

*Proof*. In a flat affine coordinate chart for the selected connection, covariant derivatives of scalar fields and their derivative tensors coincide with ordinary partial derivatives at the orders used here. Therefore *E*_*i*_ = *∂*_*i*_*µ, K*_*ij*_ = *∂*_*i*_*∂*_*j*_*µ*, and *L*_*ijk*_ = *∂*_*i*_*∂*_*j*_*∂*_*k*_*µ*.

### Proposition 2

(Coordinate covariance). *Let q and* 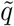 *be two coordinate systems on M. Under a coordinate change, E*_*a*_, *K*_*ab*_, *and L*_*abc*_ *transform as covariant tensors of rank one, two, and three, respectively. Therefore scalar contractions formed from these tensors and appropriate contravariant perturbation tensors are coordinate invariant*.

*Proof*. The covariant derivative of a scalar is a covector. The covariant derivative of a covector is a covariant two-tensor, and the symmetrized covariant derivative of a covariant two-tensor is a covariant three-tensor. Tensorial transformation laws follow from the definition of the connection. Complete contractions with contravariant tensors are scalars.

Table 1 summarizes the interpretation of the hierarchy.

**Table 1:**
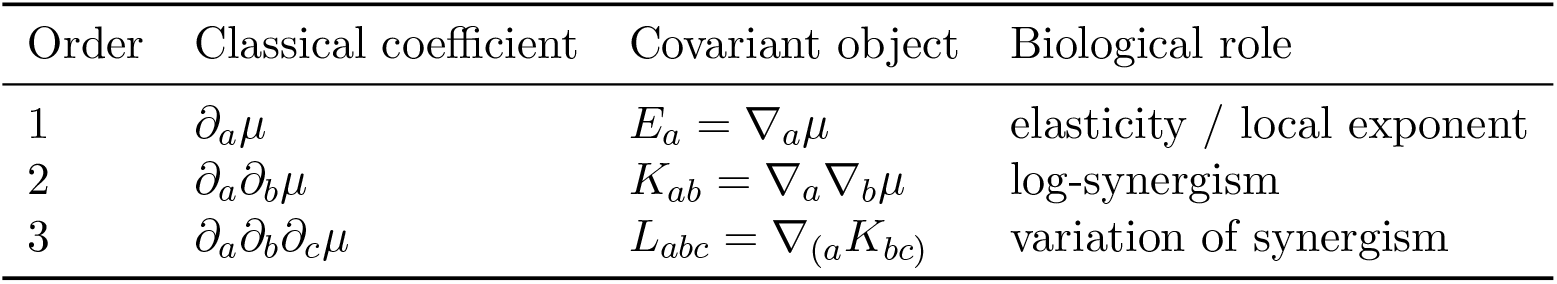
Classical and covariant interpretation of the first three BST orders. The covariant objects transform tensorially once the connection has been specified.

## 5 S-system vector fields in cBST form

An S-system writes each state equation as a difference of positive power-law fluxes,

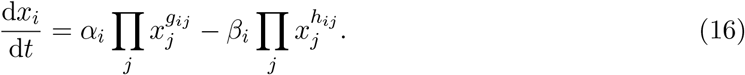

In log coordinates, the production and degradation log-fluxes are affine functions,

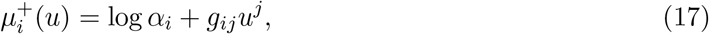

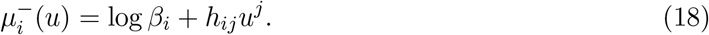

Thus exact S-system fluxes satisfy

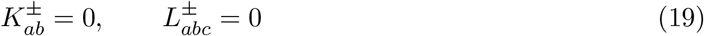

under the flat log connection. Classical S-systems are therefore exactly cBST1 models in the natural log chart.

For a general positive flux 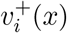or 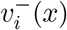, cBST provides a local hierarchy for each log-flux. cBST1 gives the local S-system exponents. cBST2 quantifies the local interaction structure that is discarded by replacing the flux by a single power law. cBST3 records how this discarded interaction changes over the operating region. In this sense, cBST does not replace S-systems; it gives a coordinate-consistent hierarchy around them.

## 6 Examples

### 6.1 Pure power law

Consider

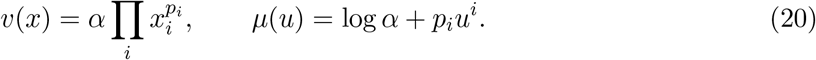

In the flat log chart,

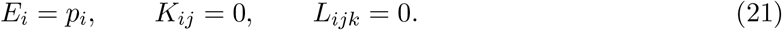

Thus an exact power law has no cBST2 or cBST3 structure. If a nonlinear coordinate transformation is introduced, the ordinary Hessian in the new coordinate system can become nonzero, but the covariant Hessian remains the tensorial transform of zero when the connection is transformed consistently. This is the simplest example showing why cBST2 should not be identified with an ordinary Hessian unless a connection is specified.

### 6.2 Michaelis–Menten saturation

For the one-substrate Michaelis–Menten form

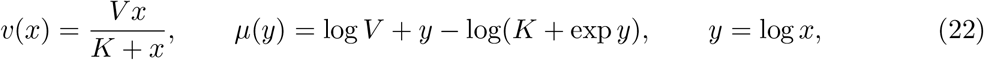

the flat-log cBST1 coefficient is

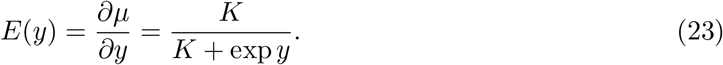

The effective local exponent decreases from one in the dilute regime to zero in the saturated regime. The cBST2 coefficient is

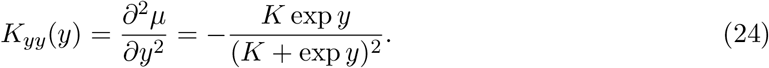

The negative sign records that the elasticity decreases as concentration increases. Its magnitude is largest near the transition between first-order and saturated regimes. This example shows how cBST2 gives a local, coordinate-consistent descriptor of saturation, not merely a formal second derivative.

### 6.3 Hill activation

For a Hill activation function

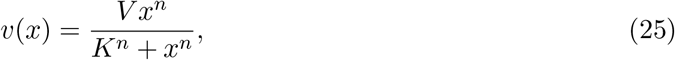

the log-response in *y* = log *x* is

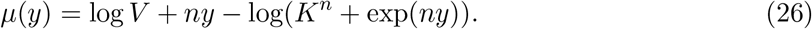

The cBST1 coefficient is

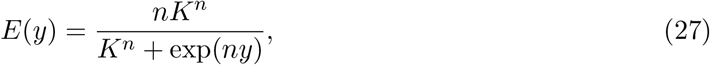

which decreases from *n* in the low-concentration regime to zero in the saturated regime. The flat-log cBST2 coefficient is

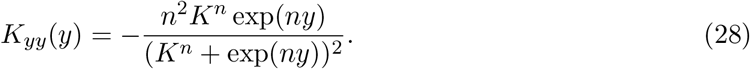

The flat-log cBST3 coefficient is

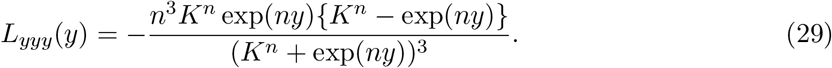

Thus the strongest second-order departure from a local power law occurs near *x* = *K*, where the response changes from cooperative power-law-like growth to saturation. The cBST3 coefficient changes sign across this transition and therefore indicates whether the magnitude of log-curvature is increasing or decreasing along the operating direction. This analytically connects a standard biochemical regulatory form to cBST1, cBST2, and cBST3.

### 6.4 Two-input log-synergism

A minimal two-input example shows why off-diagonal cBST2 components are biologically interpretable. Consider

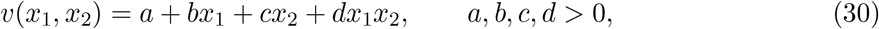

and set *u*_*i*_ = log *x*_*i*_. The log-response is

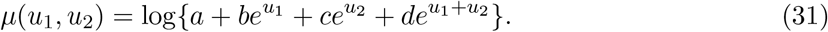

In the flat log chart, the off-diagonal cBST2 component is

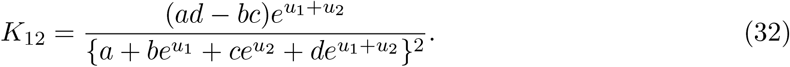

Thus *K*_12_ is positive when the joint term is stronger than expected from the two single-input terms, negative when the joint contribution is weaker, and zero when *ad* = *bc*. This gives a local differential meaning to log-synergism between two biochemical variables. It also shows why cBST2 is more than a scalar curvature diagnostic: its off-diagonal entries describe interaction structure among inputs.

### 6.5 Mixed monomial rate and exponent cumulants

Now consider a positive mixed-monomial rate,

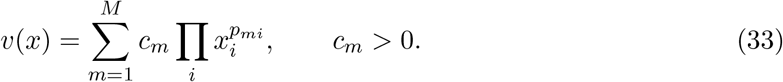

In log coordinates,

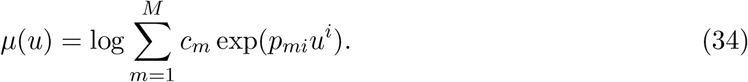

Define log-sum-exp weights

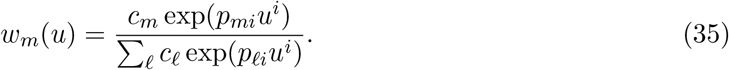

Then, in the flat log chart,

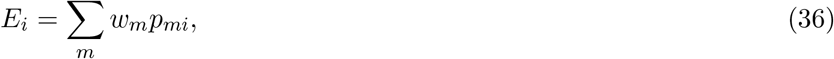

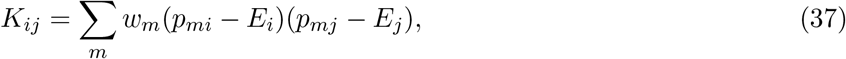

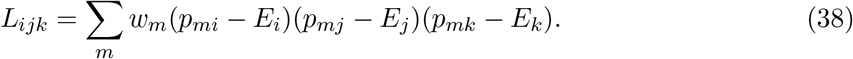

Thus the cBST tensors are cumulants of the exponent vectors *p*_*m*_ under the local weights *w*_*m*_. cBST1 is the local mean exponent, cBST2 is the local covariance of exponent vectors, and cBST3 is their local third central cumulant.

This example gives a geometric interpretation of local power-law approximation. A single dominant monomial has nearly degenerate weights, so *K* and *L* are small. When two or more monomials compete, the exponent distribution broadens and cBST2 increases. The operating point moves the weights *w*_*m*_ across the exponent set, suggesting a local connection between cBST tensors and the geometry of exponent polytopes or Newton polytopes. This connection is not required for the definitions above, but it provides intuition: cBST2 detects local mixing among exponent vectors, whereas cBST3 detects skewed transitions among competing monomial regimes.

### 6.6 A one-dimensional Schlögl-type flux

The following example is motivated by the classical Schlögl chemical reaction model for non-equilibrium phase transitions (Schlögl, 1972). The Schlögl model is widely used as a canonical bistable chemical system, and its deterministic, stochastic, and thermodynamic properties have been revisited in later work (Vellela and Qian, 2009). Here we use only its simplest mixed-monomial production structure, not the full stochastic reaction network.

For a one-dimensional autocatalytic production flux

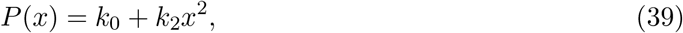

we have

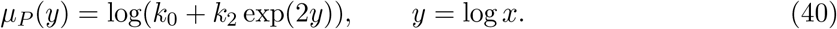

The local exponent is

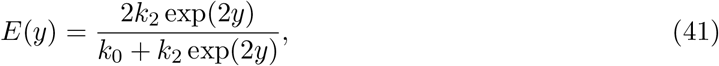

and the cBST2 curvature is

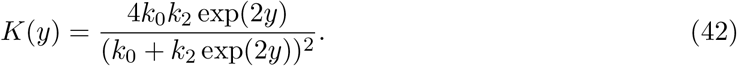

The curvature is largest in the transition region where the constant and quadratic monomials compete. This illustrates the mixed-monomial result in the smallest possible setting: local power-law behavior is reliable in a monomial-dominant regime and least reliable near a transition between monomial regimes.

## 7 Quantitative validation methods

### 7.1 Published biochemical model and operating points

The practical utility of the hierarchy was evaluated using three representative rate laws from the curated yeast glycolysis model BIOMD0000000064 in BioModels (Teusink et al., 2000; Li et al., 2010). The selected rates were glucose transport (vGLT), glucose phosphorylation (vGLK; the model’s hexokinase/glucokinase expression), and phosphofructokinase (vPFK). They provide progressively richer response structures: a two-variable transport law, a reversible four-variable enzyme rate, and a five-variable allosterically regulated rate. The rate equations and parameters were implemented from the curated model record; multiplicative compartment factors, which do not affect derivatives of the log-response, were omitted.

The reference concentrations were the model’s reported operating values under 50 mM external glucose. For vGLT, (GLC_o_, GLC_i_) = (50, 0.10). For vGLK, (GLC_i_, ATP, G6P, ADP) = (0.10, 2.51, 1.03, 1.29). For vPFK, the values were F6P = 0.11, ATP = 2.51, AMP = 0.30, F26BP = 0.020, and F16BP = 0.60. Concentrations are in mM. Table 2 summarizes the benchmark setup.

**Table 2:** Representative rate laws from BIOMD0000000064 used for quantitative validation. The dimension is the number of perturbed log-concentrations.

| Rate | Biological role | Dimension | Perturbed variables |
| --- | --- | --- | --- |
| vGLT | glucose transport | 2 | GLC<sub>o</sub>, GLC<sub>i</sub> |
| vGLK | glucose phosphorylation | 4 | GLC<sub>i</sub>, ATP, G6P, ADP |
| vPFK | phosphofructokinase | 5 | F6P, ATP, AMP, F26BP, F16BP |

**Table 3:** Prediction error and improvement at the largest tested log-perturbation radius, *ε* = 0.16. Gains are reductions relative to the preceding order.

| Rate | RMSE<sub>1</sub> | RMSE<sub>2</sub> | RMSE<sub>3</sub> | Gain<sub>2/1</sub> | Gain<sub>3/2</sub> |
| --- | --- | --- | --- | --- | --- |
| vGLT | $3.60 \times 10^{-4}$ | $1.23 \times 10^{-5}$ | $3.26 \times 10^{-7}$ | 96.6% | 97.4% |
| vGLK | $1.66 \times 10^{-3}$ | $2.99 \times 10^{-5}$ | $5.81 \times 10^{-7}$ | 98.2% | 98.1% |
| vPFK | $4.60 \times 10^{-3}$ | $6.40 \times 10^{-5}$ | $2.02 \times 10^{-5}$ | 98.6% | 68.4% |

### 7.2 Finite-perturbation benchmark

For each positive rate *v*(*x*), the exact response was *µ*(*u*) = log *v*(exp *u*) and the reference point was *u*_*\**_ = log *x*_*\**_. Log-concentration perturbations *δu* were sampled uniformly inside a Euclidean ball of radius

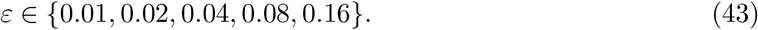

For each rate and radius, 10,000 perturbations were generated with random seed 12345. The exact response *µ*(*u*_*\**_ + *δu*) was compared with

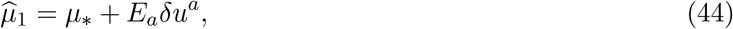

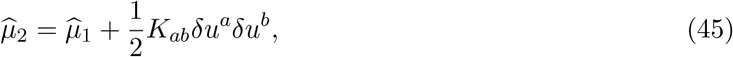

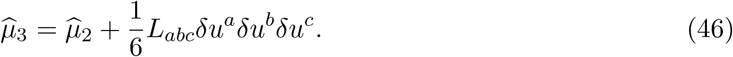

All derivative tensors were calculated in double precision by automatic differentiation. The primary outcome was the log-rate root-mean-square error

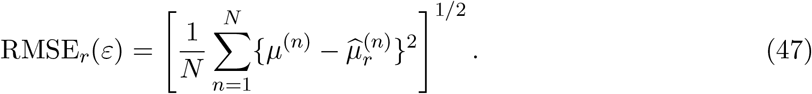

Median, 95th-percentile, and maximum absolute errors and the fraction of samples improved by each added order were also recorded.

### 7.3 Truncation-error diagnostics

To test whether the higher-order descriptors diagnose failure of lower-order approximations, the magnitudes

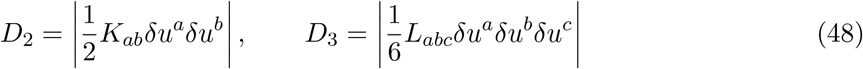

were compared with the actual cBST1 and cBST2 absolute errors, respectively. Spearman rank correlations were calculated separately for every rate and perturbation radius. This test treats cBST2 and cBST3 as local indicators of where the preceding truncation is inadequate, rather than only as descriptive coefficients.

### 7.4 Coordinate-consistency test

The flat logarithmic coordinates *u* were transformed componentwise by

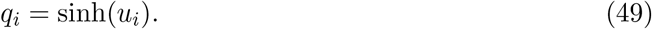

The flat connection was transformed with the chart, giving the nonzero coefficients

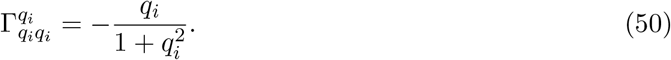

For each rate, 2,000 random unit tangent directions were generated. The scalar contractions *K*_*ab*_*ξ*^*a*^*ξ*^*b*^and*L*_*abc*_*ξ*^*a*^*ξ*^*b*^*ξ*^*c*^ were calculated in both charts after transforming the tangent components. These covariant results were compared with contractions formed from ordinary second and third partial derivatives in the nonlinear chart. Relative differences used a denominator floor of 10^*−*14^ to avoid division by numerical zero. Supplementary tests on exact power-law, Michaelis–Menten, Hill, and two-input response models verify the analytic limiting behavior and truncation orders.

## 8 Quantitative validation results

### 8.1 Order-dependent prediction accuracy

Figure 2 shows that every additional retained order reduced the finite-perturbation error for all three published-model rates. At the largest tested radius, *ε* = 0.16, the cBST1 RMSE values were 3.60 × 10^*−*4^, 1.66 × 10^*−*3^, and 4.60 × 10^*−*3^ for vGLT, vGLK, and vPFK, respectively. cBST2 reduced these errors by 96.6%, 98.2%, and 98.6%. cBST3 then reduced the remaining cBST2 errors by a further 97.4%, 98.1%, and 68.4%. The smaller additional gain for vPFK is consistent with its stronger allosteric nonlinearity and higher input dimension; even there, cBST3 improved upon cBST2 for 93.0% of the sampled perturbations at *ε* = 0.16.

**Figure 2.**
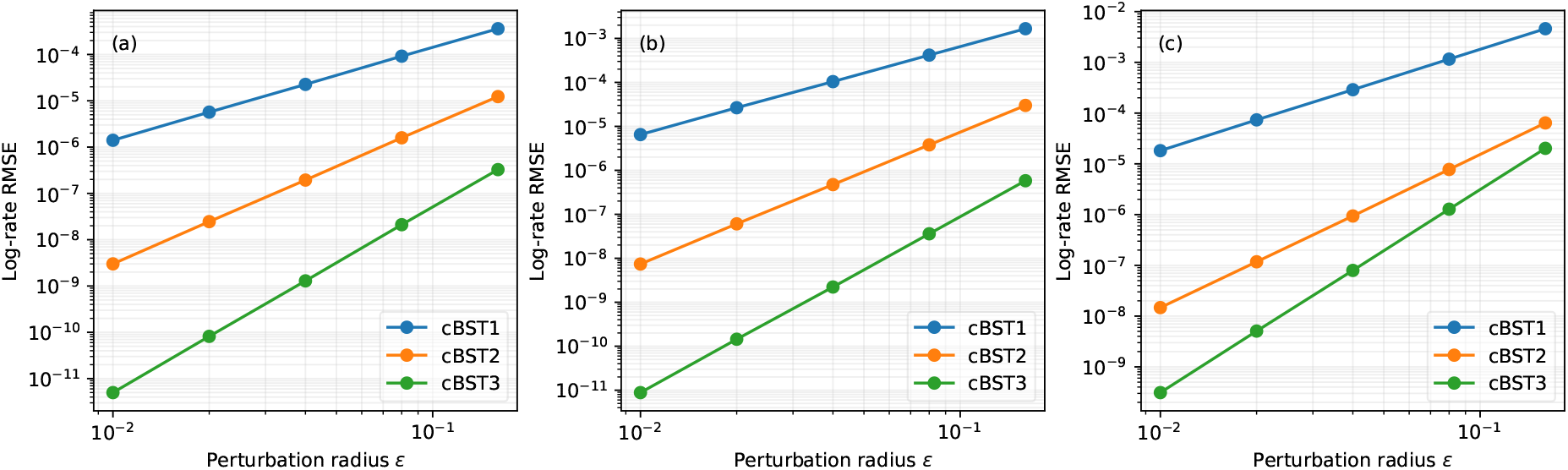
Finite-perturbation prediction errors for three representative rate laws from the curated yeast glycolysis model BIOMD0000000064: (a) vGLT, (b) vGLK, and (c) vPFK. Each point summarizes 10,000 perturbations sampled uniformly in a log-concentration ball of radius *ε*. cBST2 and cBST3 systematically reduce the error of the lower-order truncations.

### 8.2 Higher-order tensors predict lower-order failure

The quadratic contribution *D*_2_ closely tracked the actual cBST1 error. At *ε* = 0.16, the Spearman correlations were 0.9992, 0.9998, and 0.9993 for vGLT, vGLK, and vPFK. The cubic contribution *D*_3_ similarly tracked the cBST2 residual, with correlations 0.9996, 0.9986, and 0.9259. The relation remained strong across all five perturbation radii. Thus cBST2 and cBST3 provide operational diagnostics: their contractions with a proposed perturbation estimate whether the previous-order local power law is likely to be insufficient.

### 8.3 Covariant contractions remove coordinate artifacts

The coordinate test gave the same cBST contractions in *u* and *q* = sinh(*u*) to numerical precision. Across the three rates, the 95th-percentile relative mismatch was at most 4.2 × 10^*−*14^ for cBST2 and 1.2 × 10^*−*13^ for cBST3. In contrast, contractions based on ordinary partial derivatives had median relative mismatches of 0.69–1.46 at second order and 2.63–5.46 at third order. Figure 3 summarizes both the error-prediction and coordinate-consistency tests.

**Figure 3.**
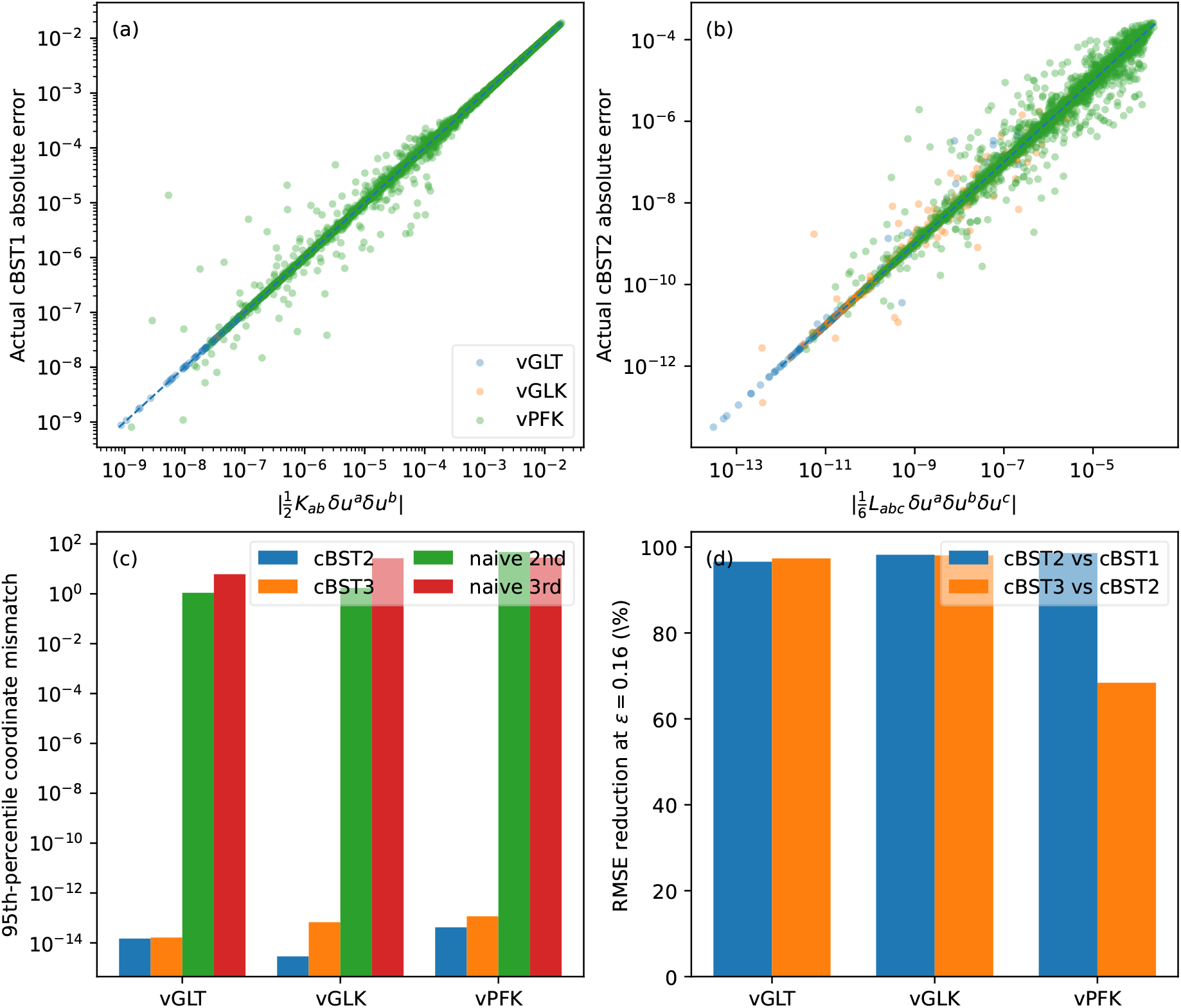
Quantitative utility and coordinate-consistency tests. (a,b) Magnitudes of the cBST2 and cBST3 correction terms versus observed errors of the preceding truncations. (c) Covariant and ordinary-derivative coordinate mismatches under *q*_*i*_ = sinh(*u*_*i*_). (d) RMSE reductions at *ε* = 0.16.

### 8.4 Analytic verification in the supplement

Supplementary Hill-response tests at an operating point away from the symmetry point recovered fitted error slopes of approximately 2, 3, and 4 for cBST1, cBST2, and cBST3, respectively, for Hill coefficients *n* = 1, 2, 3, 4. Exact power-law tests additionally confirm that a nonlinear chart can produce a nonzero ordinary Hessian while the transformed covariant Hessian remains zero. These controlled tests support the interpretation of the published-model results as the expected hierarchy of local truncation corrections rather than an artifact of one parameterization.

## 9 Interpretation and relation to approximation risk

The hierarchy separates three ideas that are often blended in local biochemical modeling. First, cBST1 describes the best local power-law exponent in the selected tangent geometry. Second, cBST2 describes the intrinsic second-order departure from a local power law. Third, cBST3 describes how that departure varies over the operating region. In a flat log chart these are simply the first three terms of the log-Taylor expansion. In nonlinear coordinates they are tensorial objects defined relative to the selected connection.

This distinction matters for comparing models across different choices of variables. A biochemical rate may be written in concentrations, normalized concentrations, logarithmic variables, reduced coordinates, saturation coordinates, or latent variables. Ordinary Hessians can change substantially across these descriptions. cBST2 changes as a tensor, not as an arbitrary collection of second derivatives. Therefore a scalar diagnostic built by contracting *K*_*ab*_ with a contravariant fluctuation covariance or perturbation tensor is coordinate invariant.

The present paper focuses on the cBST hierarchy itself. A separate approximation-risk diagnostic uses cBST2 to quantify the expected information loss caused by replacing a nonlinear log-response by its cBST1 truncation under local fluctuations (Oosawa, 2026). In that setting, the leading Gaussian risk is a scalar contraction of the covariant log-synergism tensor with the fluctuation covariance. The present hierarchy provides the geometric foundation for that risk formula, while the risk formula provides one decision-theoretic use of cBST2.

The mixed-monomial formulas clarify the relation to S-system modeling. S-system terms are exact affine functions in log space and therefore have zero cBST2 and cBST3 in the flat log chart. General rate laws can be locally represented by S-system terms through cBST1, and the discarded cBST2–cBST3 tensors quantify the local nonlinear interaction structure that the S-system term omits. This is particularly relevant to structural kinetic modeling, where local elasticities are treated as interpretable descriptors of unknown rate laws (Steuer et al., 2006; Massing et al., 2022). cBST extends this local language from first-order elasticities to coordinate-consistent higher-order descriptors.

## 10 Limitations

The covariant hierarchy requires a specified connection. There is no unique connection that is best for every biochemical modeling problem. The flat log connection is natural for classical BST and was used for the finite-perturbation benchmark. Metric-derived or information-geometric connections may be useful in statistical or observational settings, but they are optional modeling choices. Numerical values of cBST2 and cBST3 must therefore be reported together with the chosen reference connection.

The validation used local rate laws from a curated mechanistic model, not a refit of the complete glycolytic dynamics or newly collected experiments. It establishes local approximation accuracy and coordinate consistency for representative fluxes, but it does not claim improved prediction of whole-network trajectories, global steady states, identifiability, or model selection. Those questions require integration of the complete system and comparison with independent experimental observations.

The benchmarks used exact mechanistic rate expressions and double-precision automatic differentiation. Estimation of cBST2 and especially cBST3 from noisy experimental data will require regularization, uncertainty quantification, and experimental designs that excite multiple local directions. In many empirical settings, cBST2 may be the highest practically stable order. The supplied scripts and source data provide a reproducible baseline for studying those extensions. Although Eq. (14) defines a formal hierarchy beyond third order, cBST4 and higher were not included in the benchmark because the present biological questions are already resolved by elasticity, log-synergism, and its local variation. Extending the numerical study to higher ranks without an independently motivated four-way interaction or a demonstrable cBST3 residual would add parameters faster than biological interpretability.

## 11 Conclusion

A coordinate-consistent hierarchy of Biochemical Systems Theory was formulated and quantitatively evaluated. cBST1 recovers classical elasticities, cBST2 represents covariant log-synergism, and cBST3 represents its local covariant variation. Tests on three representative rates from a curated yeast glycolysis model showed systematic order-by-order reductions in finite-perturbation prediction error. The cBST2 and cBST3 correction terms also predicted where lower-order approximations failed, turning the hierarchy into a practical local model-order diagnostic. Nonlinear reparameterization tests confirmed that covariant contractions preserve the response description while ordinary higher derivatives acquire substantial coordinate artifacts. Together with the supplementary analytic verification and reproducible source data, these results support cBST as a bridge between classical power-law modeling, structural kinetic interpretation, and quantitative approximation-risk assessment.

## Author contributions

Chikoo Oosawa: Conceptualization, Methodology, Formal analysis, Software, Writing—original draft, Writing—review and editing.

## Data and code availability

This study generated quantitative benchmark data by evaluating representative rate laws from the curated yeast glycolysis model BIOMD0000000064 and analytically controlled response models across operating points, perturbation amplitudes, and nonlinear coordinate representations. All processed CSV outputs, figure source data, environment metadata, and Python scripts required to reproduce the analyses are included in the accompanying package. The original model is available from BioModels under accession BIOMD0000000064. No newly collected experimental data were used.

## Declaration of generative AI and AI-assisted technologies

During the preparation of this work, the author used ChatGPT to assist with language polishing, structural organization, and LaTeX package preparation. After using this tool, the author reviewed and edited the content as needed and takes full responsibility for the manuscript.

## A Coordinate artifact in a one-dimensional example

Let *µ*(*u*) = *au* + *b* be an exact log-affine response. In the flat *u* coordinate, *K*_*uu*_ = 0. Now set *u* = *φ*(*q*) with nonlinear *φ*. The ordinary second derivative in *q* is

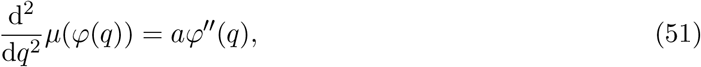

which is generally nonzero. This is not intrinsic log-synergism; it is produced by the coordinate transformation. The transformed flat connection has

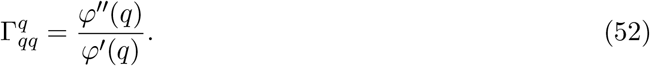

Using Eq. (10), the covariant Hessian in *q* is

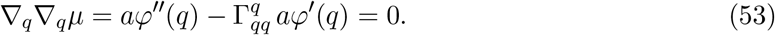

Thus cBST2 correctly reports that an exact power-law response remains exact after reparameterization.

## B Remarks on curvature and third derivatives

For a scalar *µ* and a torsion-free connection, the second covariant derivative is symmetric. At third order, commutators of covariant derivatives can introduce curvature terms when acting on the covector ∇*µ*. The symmetrized definition in Eq. (13) is therefore used as the coordinate-consistent local cubic coefficient in the geodesic Taylor expansion. In a flat log chart, all such curvature issues disappear and cBST3 is the ordinary third derivative.

